# Markerless motion capture estimates of lower extremity kinematics and kinetics are comparable to marker-based across 8 movements

**DOI:** 10.1101/2023.02.21.526496

**Authors:** Ke Song, Todd J. Hullfish, Rodrigo Scattone Silva, Karin Grävare Silbernagel, Josh R. Baxter

## Abstract

Motion analysis is essential for assessing in-vivo human biomechanics. Marker-based motion capture is the standard to analyze human motion, but the inherent inaccuracy and practical challenges limit its utility in large-scale and real-world applications. Markerless motion capture has shown promise to overcome these practical barriers. However, its fidelity in quantifying joint kinematics and kinetics has not been verified across multiple common human movements. In this study, we concurrently captured marker-based and markerless motion data on 10 healthy subjects performing 8 daily living and exercise movements. We calculated the correlation (R_xy_) and root-mean-square difference (RMSD) between markerless and marker-based estimates of ankle dorsi-plantarflexion, knee flexion, and three-dimensional hip kinematics (angles) and kinetics (moments) during each movement. Estimates from markerless motion capture matched closely with marker-based in ankle and knee joint angles (R_xy_ ≥ 0.877, RMSD ≤ 5.9°) and moments (R_xy_ ≥ 0.934, RMSD ≤ 2.66 % height × weight). High outcome comparability means the practical benefits of markerless motion capture can simplify experiments and facilitate large-scale analyses. Hip angles and moments demonstrated more differences between the two systems (RMSD: 6.7° – 15.9° and up to 7.15 % height × weight), especially during rapid movements such as running. Markerless motion capture appears to improve the accuracy of hip-related measures, yet more research is needed for validation. We encourage the biomechanics community to continue verifying, validating, and establishing best practices for markerless motion capture, which holds exciting potential to advance collaborative biomechanical research and expand real-world assessments needed for clinical translation.

## 1. Introduction

Motion analysis is essential for assessing in-vivo human biomechanics. The two main components of motion analysis are kinematics and kinetics. Kinematics quantify the position, orientation, and movement of human body segments and joints, while kinetics characterize the mechanical forces and moments needed to produce these movements (Cappozzo et al., 2005; Robertson et al., 2013). Accurately quantifying human biomechanics relies on experimental and computational methods to capture and track the dynamic positions of body segments. The validity of a motion capture system thus depends on its accuracy and consistency to estimate kinematic and kinetic parameters, including joint angles and moments during common movements of daily living or exercises (Camomilla et al., 2017; Cappozzo et al., 2005).

Marker-based motion capture is the standard to analyze human motion. This approach typically tracks reflective markers attached on skin or clothes using infrared cameras to estimate the three-dimensional (3D) kinematic motion of bony segments. Kinematics are then coupled with external force measurements to compute kinetics (Camomilla et al., 2017; Robertson et al., 2013). These computations rely on marker-based approximation of bony landmarks, which are inherently inaccurate due to marker misplacement and skin motion relative to the bones (Della Croce et al., 2005; Leardini et al., 2005). Marker-based systems are also expensive, require extensive experimental setup, and time-intensive data processing that can introduce errors (Chiari et al., 2005). These factors all limit its utility in large-scale applications, especially in clinical and real-world settings where such a system is not accessible, affordable, or practical. There is a clear need to develop techniques that improve the accuracy, consistency, and feasibility of motion capture to better assess biomechanics in clinically relevant environments.

Markerless motion capture developed in recent years has shown promise to overcome the practical barriers of marker-based systems. This new technique uses computer vision and machine learning to directly detect human body landmarks from digital images (Colyer et al., 2018; Cronin, 2021; Drazan et al., 2021; Kanko et al., 2021a; Mathis et al., 2018) instead of relying on specialized cameras to track physical markers. Markerless systems take advantage of powerful algorithms and automated computation that eliminates many manual processing steps and error sources inherent to marker-based techniques. Recent studies have shown that markerless technique improves inter-session repeatability (Kanko et al., 2021b) and is robust against subject variations such as clothing (Keller et al, 2022). Markerless motion capture is potentially better suited for analysis in real-world environments because digital images can also be recorded using consumer-grade devices such as smartphone cameras. However, the fidelity of markerless motion capture in quantifying 3D joint kinematics and kinetics has not been fully verified against standard marker-based techniques. To our knowledge, only three recent studies compared lower extremity 3D kinematics or kinetics from concurrent marker-based and markerless motion capture (Ito et al., 2022; Kanko et al., 2021a; Tang et al., 2022). Kanko et al. (2021a) found comparable kinematics between the two systems during treadmill walking. Ito et al. (2022) added that kinematics during squatting and forward hopping were also comparable in the sagittal plane, but not in frontal and transverse planes. The only kinetic study to date is by Tang et al. (2022), which reported between-system differences in peak joint moments and powers during treadmill running. These studies either only assessed kinematics (Ito et al., 2022; Kanko et al., 2021a) or a single movement (Kanko et al., 2021a; Tang et al., 2022). No study has compared lower extremity 3D kinematics and kinetics between marker-based and markerless systems across multiple daily living or exercise movements.

In this study, we compared lower extremity kinematics and kinetics estimated from marker-based and markerless motion capture across 8 daily living and exercise movements. We hypothesized that estimates from markerless motion capture would match closely with marker-based system at the ankle and knee joints during most movements, while differences between the two systems would be larger at the hip joint and during faster movements due to limitations from both techniques.

## 2. Methods

### 2.1. Subjects

We recruited 10 healthy adult subjects (5M/5F, 21.9 ± 1.9 years old, body mass index = 23.8 ± 2.4 kg/m^2^) from our University campus and local community. Subjects were between 18 and 40 years of age and had no self-reported history of injury in the lower limbs or spine in the last 6 months. An experienced physical therapist (R.S.S.) confirmed that each subject had no current injury in the lower limbs or lower back which could interfere with their movement patterns. Subjects provided written informed consent before participating and the study was approved by the University of Pennsylvania’s Institutional Review Board.

### 2.2. Motion capture systems and experiment

We positioned twelve near-infrared marker-based cameras (1.3-megapixel Raptor-E ×10, 12-megapixel Raptor-12 ×2, Motion Analysis Corp., Rohnert Park, CA) and eight high-resolution video cameras (2.1-megapixel Optitrack Prime Color, NaturalPoint Inc., Corvallis, OR) evenly spaced around the center of our laboratory space approximately 3 meters above the floor. We used a synchronization module for concurrent motion capture of the same trials, with the marker-based system providing a signal to automatically start and stop the markerless camera recordings. Both systems recorded at 100 Hz. Three force platforms (BP600900, AMTI, Watertown, MA) embedded under the floor at the center of capture space recorded ground reaction forces at 1000 Hz and were synchronized to both marker-based and markerless systems.

Subjects wore exercise clothing (running shorts and tank tops) and standardized running shoes. A single experienced examiner (K.S.) placed 31 retro-reflective skin markers over anatomic landmarks of the pelvis, both thighs, shanks, feet (on the shoes) and torso to track body segment positions (Slater et al., 2018) (**Figure 1A**). We affixed markers on the posterior pelvis using a rigid cluster plate and secured it with elastic bands around the waist to ensure its visibility by the marker-based cameras. We captured a static trial with subject standing in an anatomical pose and then removed 4 calibration-only markers on medial knees and ankles so they would not interfere with each subject’s natural movements.

**Figure 1.**
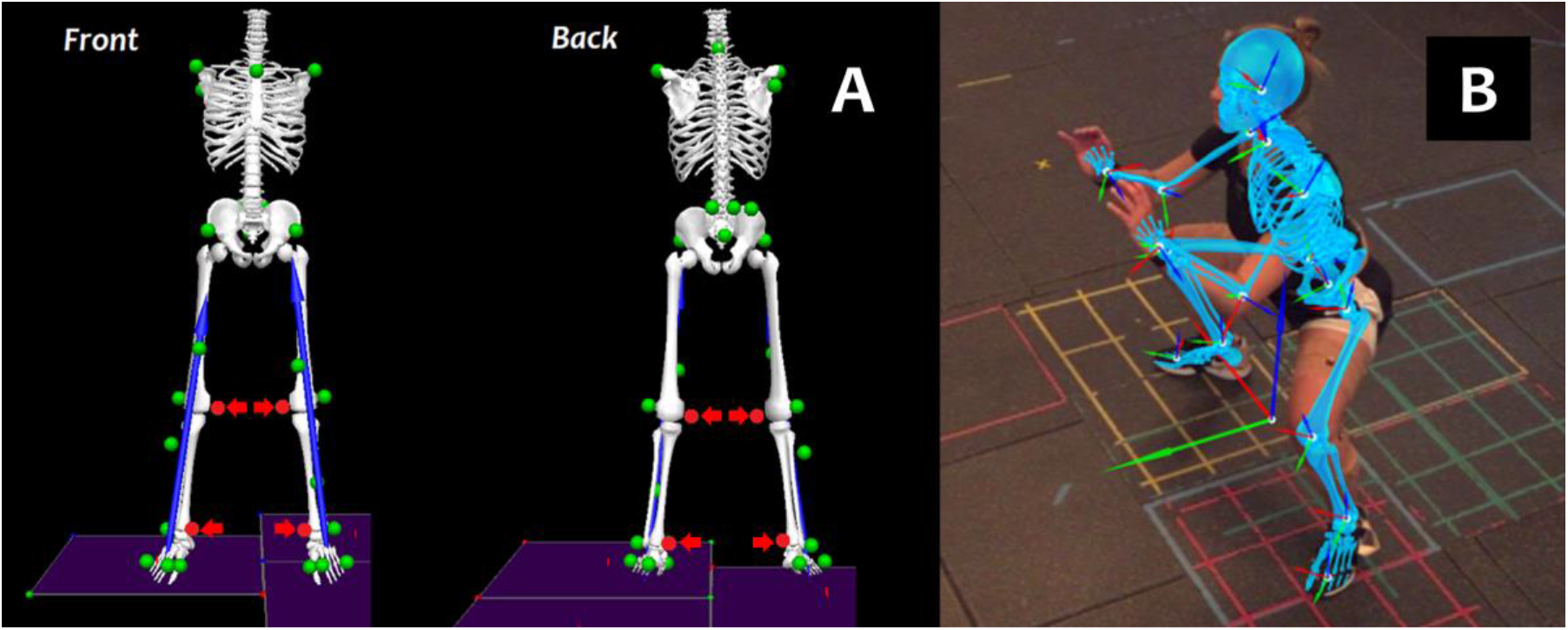
**(A)** Marker set to track the positions of pelvis, lower extremities, and torso during marker-based motion capture. We removed calibration-only markers (arrows) after an anatomical pose trial. **(B)** Example video frame of markerless motion capture overlaid with the built-in model that tracks the motion of body segments. The subject was performing a wide-stance sumo squat in this example.

Each subject performed eight (8) movement trials that are either functional activities of daily living or common exercises in sports and recreation: heel raises, over-ground walking, stepping down a 20-cm box, over-ground running, regular double-leg squat, wide-stance double-leg squat (“sumo squat”, **Figure 1B**), countermovement jump, and a run-and-cut maneuver. All subjects performed the 8 movements in this specific order and we provided 3 to 5-minute rest periods between trials to prevent physical exhaustion. They were provided visual demonstrations, practiced each movement, and received verbal guidance by a physical therapist (R.S.S.) throughout each trial. We collected at least 3 successful repetitions of each movement. These 8 movements were also part of a larger study in which we captured the movements of 36 exercises or daily living activities. While we focused on the 8 representative movements in this study, we also extended our biomechanical analysis to the other 28 movements and included them in **Supplementary Material 1**.

### 2.3. Joint angles and moments

For marker-based motion capture, we manually labeled the marker trajectories to ensure they continuously tracked the correct positions of the pelvis and lower body segments. We filled small marker trajectory gaps (< 0.3 second) using cubic splines and assigned labeled trajectories to a biomechanical model (Visual3D, C-Motion, Germantown, MD). We used a link-segment model with inverse kinematics constraints to solve for the best estimates of segment positions (Robertson et al., 2013; Slater et al., 2018). This model constrained the knee joint to 1 degree of freedom (flexion-extension only), while allowing the hip and ankle joints for full 3D rotations. Slater et al. (2018) demonstrated that this model can track lower extremity segments with high fidelity with the simple marker set we used.

For markerless motion capture, we processed the 8 synchronized high-resolution videos of each trial using a commercially available software (Theia3D, Theia Markerless Inc., Kingston, ON, Canada). This software detects and estimates 3D human salient features based on a deep learning algorithm trained on over 500,000 annotated human images (Kanko et al., 2021a, 2021b, 2021c). It then scales a link-segment model to fit the subject-specific landmarks and uses inverse kinematics to solve for segment positions similar to the marker-based algorithm (Robertson et al., 2013). The built-in markerless model allows for 3 rotational degrees of freedom at the hip, knee, and ankle joints.

We calculated lower extremity joint angles during each trial from both marker-based and markerless models on the right leg of each subject as the representative side. We computed five (5) joint angles as our primary kinematic measures: ankle dorsi-plantarflexion, knee flexion, hip flexion-extension, hip adduction-abduction, and hip internal-external rotation. We focused our analysis on knee and ankle biomechanics to the sagittal plane because past studies found off-sagittal knee motion to be highly susceptible to skin artifacts (Akbarshahi et al., 2010), and high-fidelity hindfoot kinematics require capturing barefoot motion (Nester et al., 2007) which is impractical for analyzing many movements.

We calculated lower extremity joint moments from both models using the same inverse dynamics algorithm (Robertson et al., 2013). We confirmed that ground reaction forces were correctly assigned to the corresponding feet in both models, including through the box during the step-down movement. We computed ankle plantar-dorsiflexion, knee extension, hip extension-flexion, hip abduction-adduction, and hip external-internal rotation moments as our five kinetic measures, which matched the five kinematic measures. As a secondary analysis, we also calculated and compared ankle, knee, and hip joint center locations tracked by the marker-based and markerless systems as a partial contributor to joint kinetics (Robertson et al., 2013). Data from the joint center location analysis are included in **Supplementary Material 2**.

### 2.4. Data analysis

We computed joint angles and moments during individual repetitions within each captured movement trial. We identified the repetitions using ground reaction forces or manually defined start and stop events that represent the weight-bearing phases of each movement (e.g., the stance phase of walking). We normalized joint moments by subject body height times weight (Andriacchi and Strickland, 1985; Moisio et al., 2003). We computed Pearson correlation coefficients (Rxy) between markerless and marker-based estimates of each measure to quantify the agreement of the two waveforms. We also computed root-mean-square differences (RMSD) between the two estimates of each measure to quantify their average magnitude difference over each individual repetition.

We averaged each R_xy_ and RMSD across repetitions within a trial during each movement. We then calculated their group mean across the 10 subjects to determine an overall level of agreement (R_xy_) and magnitude difference (RMSD) between markerless and marker-based estimates. According to the guidelines by Schober et al. (2018), we defined that a coefficient of R_xy_ ≥ 0.7 suggests a strong correlation between the two systems, and R_xy_ ≥ 0.9 suggests a very strong correlation. In absence of a research community consensus, we defined that the between-system magnitude difference is minimal if RMSD ≤ 5° for joint angles (Akbarshahi et al., 2010; Leardini et al., 2005), and ≤ 2.5 % height (H) × weight (W) for joint moments.

## 3. Results

### 3.1. Kinematics: joint angles

Kinematic estimates from markerless motion capture matched closely to marker-based at the ankle and knee joints, while hip angles demonstrated less agreement (**Figure 2, 3**). Ankle dorsi-plantarflexion showed very strong correlations between the two systems in all movements (R_xy_ ≥ 0.900) and had consistently small RMSD (≤ 5.9°) (**Figure 2A, 3A**). Knee flexion had strong correlations for all but one movement (R_xy_ ≥ 0.877), the heel raises which involve minimal knee or hip motion (**Figure 2B, 3B**). Knee flexion RMSD were small for all movements including the heel raises (≤ 5.9°). Hip flexion had very strong correlations in all movements except the heel raises (R_xy_ ≥ 0.965). However, we observed a consistent offset where markerless motion capture estimated smaller hip flexion (RMSD: 6.7° – 13.8°) (**Figure 2C, 3C**). Hip adduction-abduction met our a priori threshold for a strong correlation (R_xy_ ≥ 0.739), but the coefficients were smaller than the sagittal joint angles. Hip adduction-abduction RMSD remained small and comparable to ankle dorsi-plantarflexion and knee flexion (≤ 5.1°; **Figure 2D, 3D**). Hip internal-external rotations showed weaker correlations between the systems (R_xy_ ≤ 0.601 for all except sumo squat; **Figure 2E, 3E**). The markerless estimates also demonstrated an offset towards hip external rotation compared to marker-based, which resulted in larger RMSD (8.2° – 15.9°). Across the 8 movements, running and run-and-cut showed the least strong correlations between the two systems, and RMSD were among the highest (**Figure 2d, 3h**).

**Figure 2.**
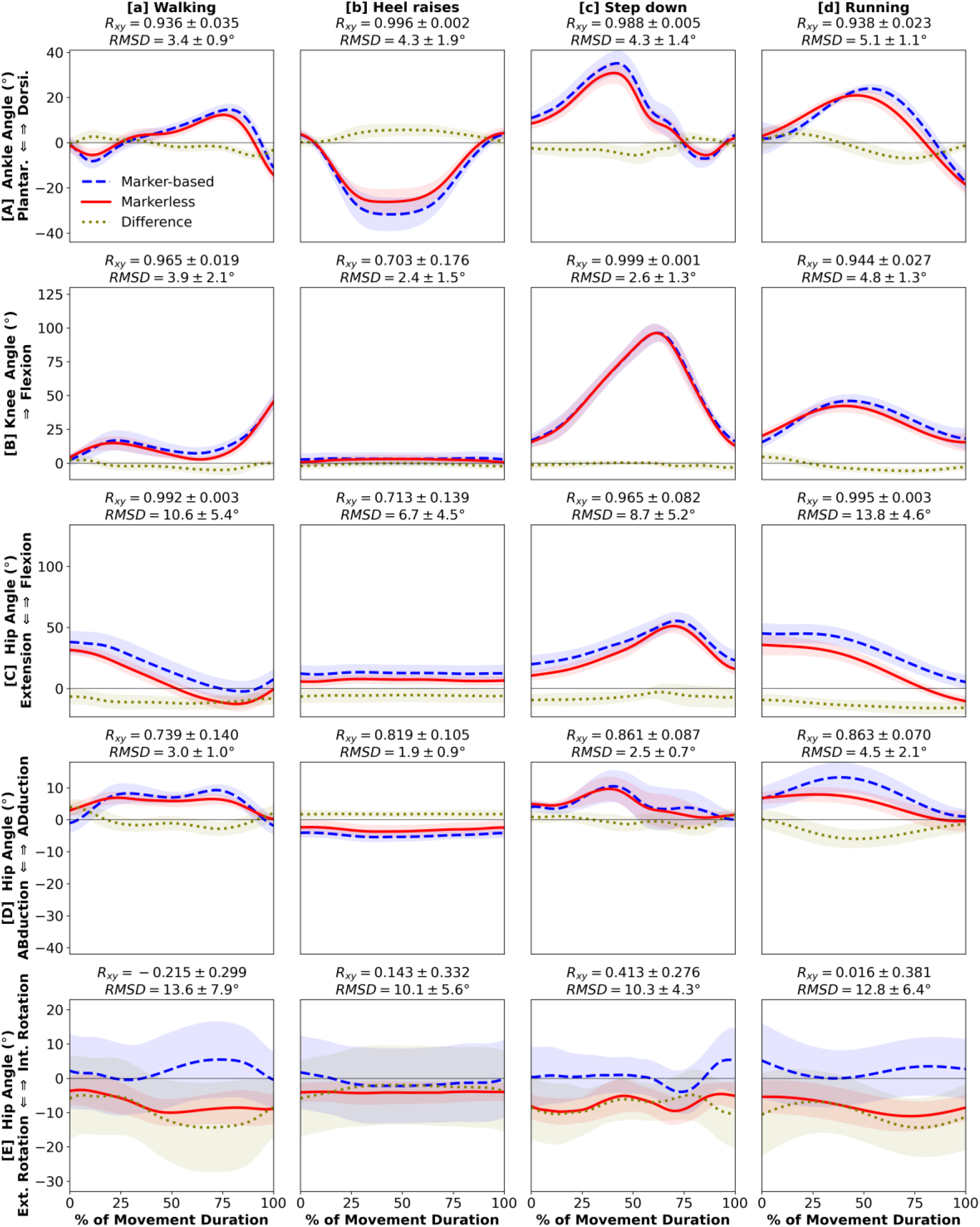
Kinematics: [**A**] ankle dorsiflexion (+) / plantarflexion (-), [**B**] knee flexion (+), [**C**] hip flexion (+) / extension (-), [**D**] hip adduction (+) / abduction (-), [**E**] hip internal (+) / external (-) rotation angles during 4 daily living movements: [**a**] walking, [**b**] heel raises, [**c**] step-down, [**d**] running. Waveforms = group mean (line) ± 1 SD (shade) for marker-based (blue), markerless (red), and their difference (gold).

**Figure 3.**
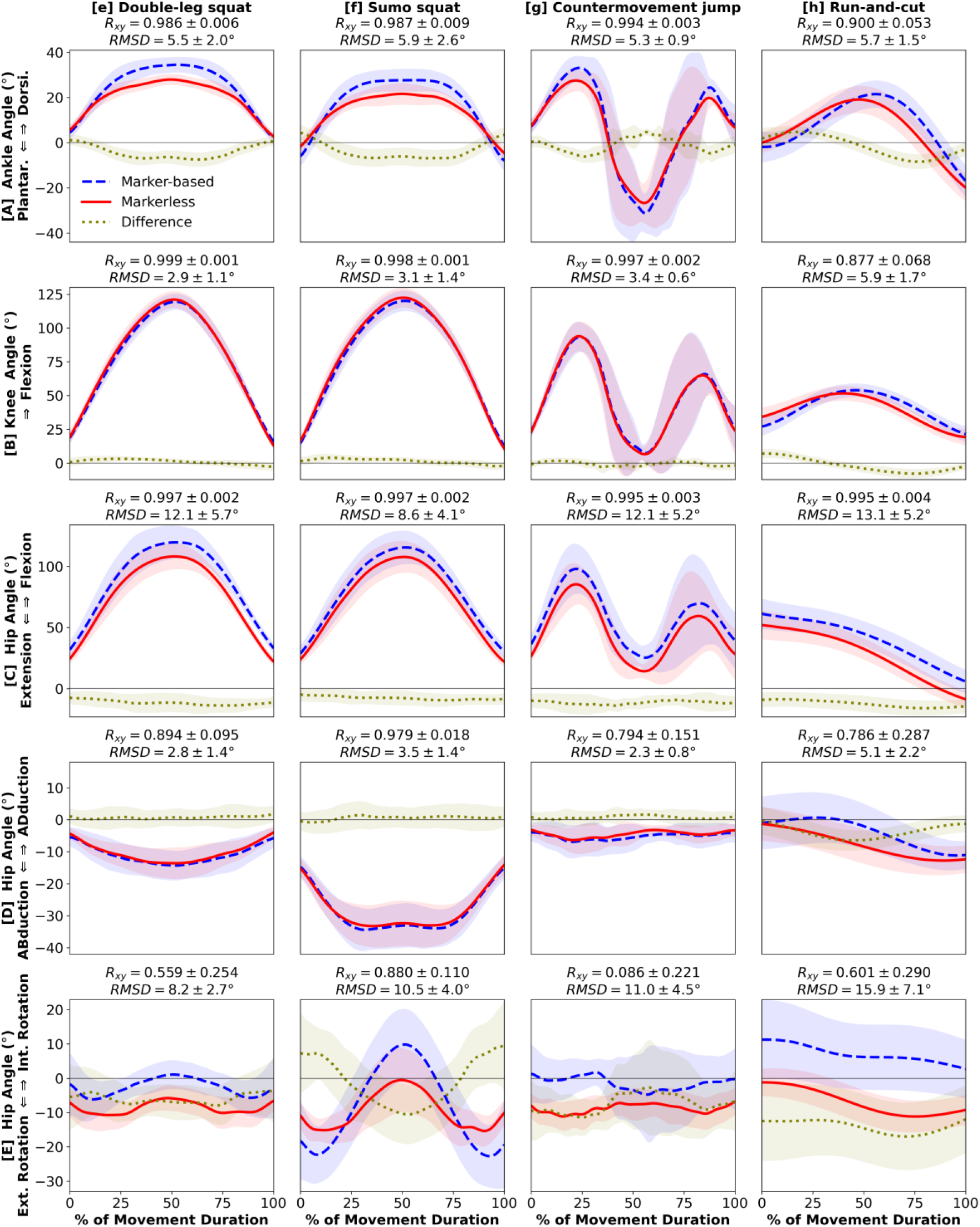
Kinematics: [**A**] ankle dorsiflexion (+) / plantarflexion (-), [**B**] knee flexion (+), [**C**] hip flexion (+) / extension (-), [**D**] hip adduction (+) / abduction (-), [**E**] hip internal (+) / external (-) rotation angles during 4 exercise movements: [**e**] double-leg squat, [**f**] sumo squat, [**g**] countermovement jump, [**h**] run-and-cut. Waveforms = group mean ± 1 SD.

### 3.2. Kinetics: joint moments

Most kinetic estimates from markerless motion capture demonstrated strong waveform agreement and magnitude similarity to marker-based (**Figure 4, 5**). We found very strong between-system correlations and low RMSD for both ankle plantar-dorsiflexion moment (R_xy_ ≥ 0.982, RMSD ≤ 1.44 % H×W; **Figure 4A, 5A**) and knee extension moment (R_xy_ ≥ 0.934, RMSD ≤ 2.66 % H×W; **Figure 4B, 5B**). Hip extension moment had strong correlations for all 8 movements (R_xy_ ≥ 0.785) and low RMSD for 6 of 8 (RMSD ≤ 1.35 % H×W; **Figure 4C, 5C**), except running (RMSD = 4.46 % H×W, **Figure 4d**) and run-and-cut (RMSD = 7.15 % H×W, **Figure 5h**) which had larger differences between markerless and marker-based estimates. Finally, we observed strong correlations and low RMSD across all 8 movements for hip abduction-adduction moment (R_xy_ ≥ 0.819, RMSD ≤ 1.42 % H×W; **Figure 4D, 5D**) and external-internal rotation moment (R_xy_ ≥ 0.821, RMSD ≤ 0.51 % H×W; **Figure 4E, 5E**). In general, running and run-and-cut showed the largest between-system kinetic differences across the 8 movements (**Figure 4d, 5h**), which matched the kinematic results.

**Figure 4.**
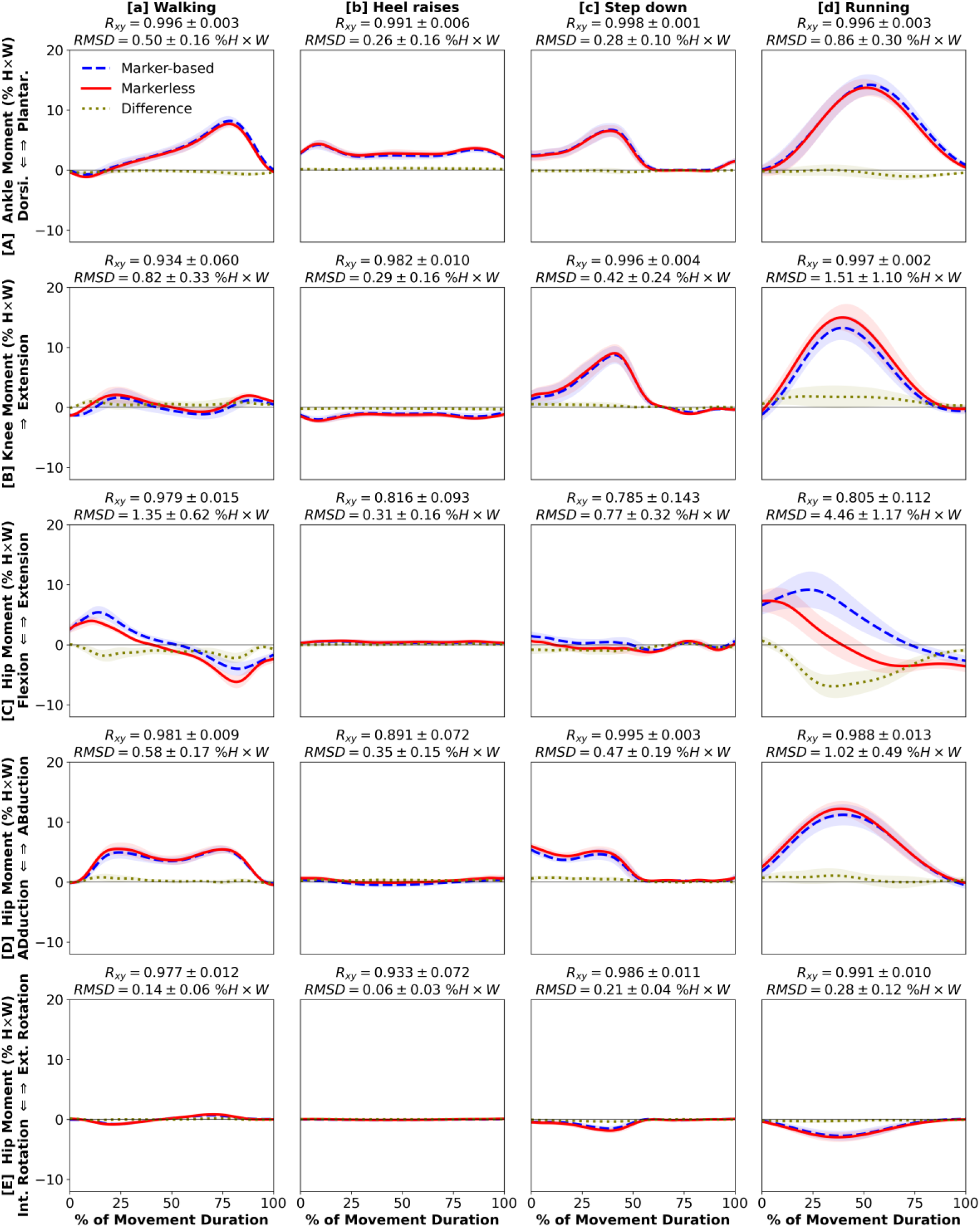
Kinetics: [**A**] ankle plantarflexion (+) / dorsiflexion (-), [**B**] knee extension (+), [**C**] hip extension (+) / flexion (-), [**D**] hip abduction (+) / adduction (-), [**E**] hip external (+) / internal (-) rotation moments during 4 daily living movements: [**a**] walking, [**b**] heel raises, [**c**] step-down, [**d**] running. Waveforms = group mean ± 1 SD.

**Figure 5.**
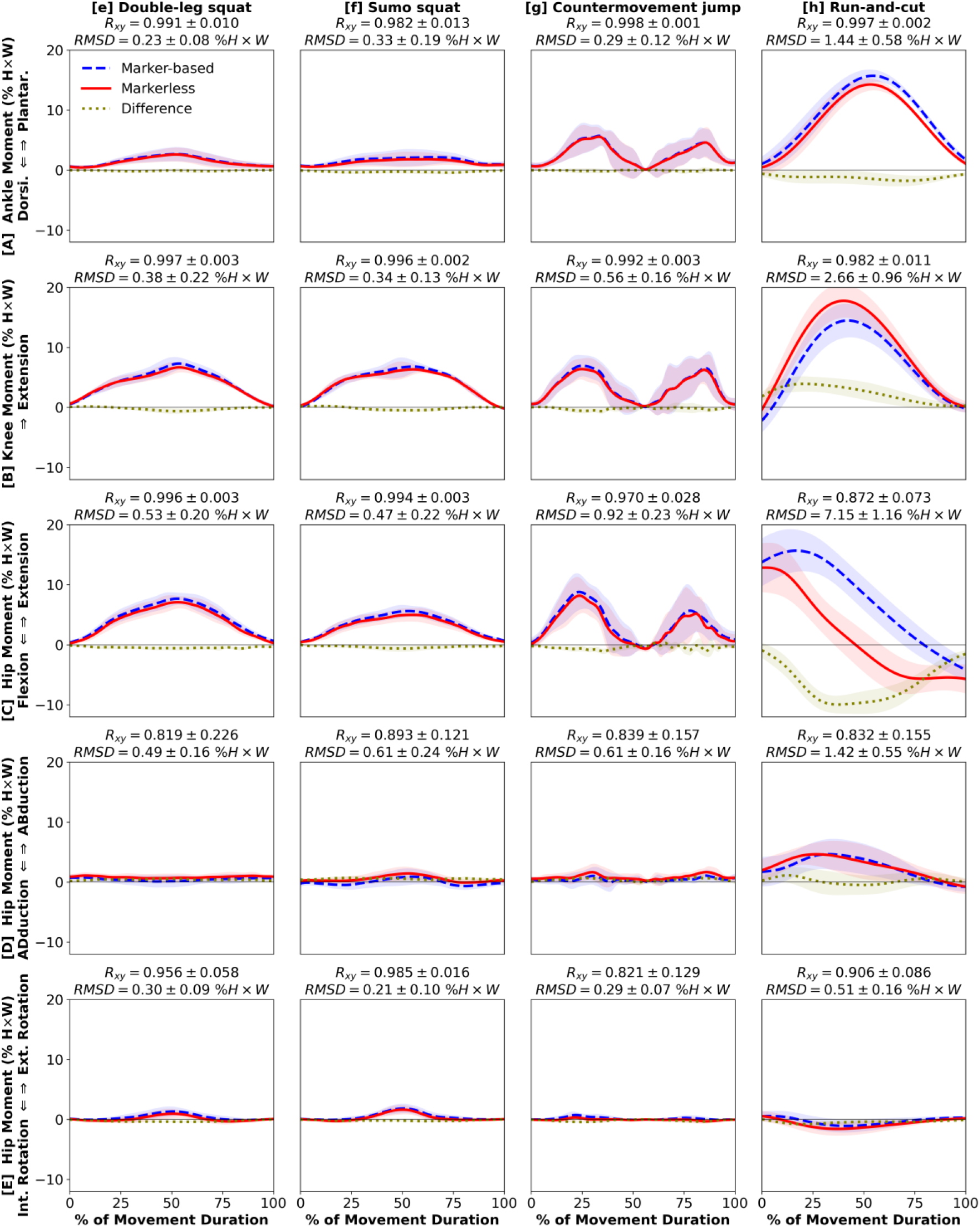
Kinetics: [**A**] ankle plantarflexion (+) / dorsiflexion (-), [**B**] knee extension (+), [**C**] hip extension (+) / flexion (-), [**D**] hip abduction (+) / adduction (-), [**E**] hip external (+) / internal (-) rotation moments during 4 exercise movements: [**e**] double-leg squat, [**f**] sumo squat, [**g**] countermovement jump, [**h**] run-and-cut. Waveforms = group mean ± 1 SD.

## 4. Discussion

We compared lower extremity kinematics and kinetics estimated from marker-based and markerless motion capture across 8 daily living and exercise movements. We confirmed our hypothesis that estimates from markerless motion capture would match closely with marker-based at the ankle and knee during most movements, while between-system differences would be larger at the hip and during faster movements.

Our results support initial investigations by Kanko et al. (2021a) that lower extremity kinematics are highly comparable between marker-based and markerless motion capture during walking. We also confirmed that the tracking quality of markerless motion capture during over-ground walking is similar to walking on treadmill (Kanko et al., 2021a). We found that the strong between-system correlations held true for the 3 sagittal lower extremity joint angles during all 8 movements we analyzed (**Figure 2A–C, 3A–C**). Waveform correlations were less strong in hip adduction-abduction and internal-external rotation (**Figure 2DE, 3DE**), possibly due to minimal off-sagittal hip motion in the 8 movements we analyzed. For most kinematic measures, the magnitude differences were small, albeit offsets in hip flexion-extension and internal-external rotation (**Figure 2CE, 3CE**). These consistent offsets are likely a result of distinct definitions of pelvis orientation (Kanko et al., 2021a) and not a consequence of kinematic tracking quality. Therefore, we conclude that the accuracy of lower extremity kinematics is generally comparable between marker-based and markerless systems. Our results are inconclusive regarding the accuracy of hip internal-external rotation. Further studies should analyze movements requiring substantial hip transverse rotations and use a marker-based model that matches the pelvis definition of the markerless model to verify whether the two systems estimate comparable values.

Our study is the first to show that, across multiple movements, lower extremity kinetics estimated from markerless and marker-based motion capture match closely on both waveform and magnitude. To date, only one study compared 3D joint kinetics from markerless system to marker-based (Tang et al., 2022). This study reported that markerless system estimated a higher peak hip flexion moment, lower knee extension moment, and higher ankle plantarflexion moment during the weight-bearing phase of treadmill running (Tang et al., 2022). Our findings did not fully agree with these observations, as our markerless system estimated a lower hip extension moment, slightly higher peak knee extension moment, and comparable ankle plantarflexion moment during over-ground running (**Figure 4d**). Several factors could have contributed to these disagreements, including distinct over-ground and treadmill running kinetics (Riley et al., 2008), moment normalization techniques (Moisio et al., 2003), marker placements, and marker-based model definitions. It is possible that the rapid running motion negatively affected the accuracy of both systems. We found relatively small magnitude differences in all kinetic measures across all movements except running and run-and-cut (RMSD ≤ 1.35 % H×W; **Figure 4, 5**). Therefore, we conclude that for slower-speed movements, markerless motion capture estimates of lower extremity kinetics are highly comparable to marker-based.

The larger between-system differences during faster movements can be explained by limitations from both techniques. Marker-induced kinematic errors from increased skin-to-bone motion are magnified when used to compute kinetics (Camomilla et al., 2017), which may explain why hip extension moments showed larger disagreements than flexion angles during running and run-and-cut. Between-system differences can also be contributed from markerless motion capture due to video image blurs during rapid movements (Kanko et al., 2021a). However, unlike the inherent skin-to-bone motion, video clarity can be optimized by adjusting system parameters such as camera resolution, lighting, shutter speed, and capture rate. We considered all these factors when setting up our markerless motion capture system.

The hip is the most error-prone lower extremity joint in marker-based biomechanical analysis. Larger errors are expected due to substantial skin-to-bone motion and the practical challenges in identifying anatomical landmarks and securing markers on the pelvis (Della Croce et al., 2005; Leardini et al., 2005; Stagni et al., 2000). During the late weight-bearing phases of running and run-and-cut, the hip must be supplied with substantial flexion moments to execute push-off (Hamner et al., 2010; Novacheck, 1998). In our results, this biomechanical phenomenon is better characterized by the markerless system (**Figure 4d, 5h**). We thus speculate that markerless motion capture estimated hip kinetics more accurately than our marker-based system. However, the absolute accuracy of hip measures is beyond the scope of our analysis without knowing the true movements of the underlying hip bones. Future studies should verify whether markerless motion capture is indeed more accurate than marker-based by comparing both motion capture techniques to concurrent assessment of hip articular dynamics, using gold-standard measurement techniques such as the bi-planar fluoroscopy (Kapron et al., 2014; Li et al., 2004; Tashman and Anderst, 2003).

Our results should be interpreted with several limitations. One important limitation is the differences in biomechanical models between the marker-based and markerless systems. The comparability of motion analyses is often hindered by different model definitions such as pelvic tilt and neutral ankle angles (Wu et al., 2002). This is evident in our data, as the hip angle offsets (**Figure 2CE, 3CE**) seem to be caused by different pelvis segment definitions. While model definition standards have been proposed (Derrick et al., 2020; Wu et al., 2005, 2002), researchers often had to make project-specific modifications due to the practical constraints of marker-based experiments. Markerless motion capture helps overcome this barrier by implementing consistent segment and joint models across multiple experiments, eliminating this source of kinematic offsets. We encourage researchers to cross-validate outcomes from markerless motion capture studies and establish best practices for yielding meaningful data. We believe such efforts would significantly advance the collaborative research and comparable clinical assessments of human biomechanics. A second limitation is that out study focused on the joint kinematics and kinetics of the lower extremity. A past study has compared upper extremity kinematics between marker-based and markerless systems (Kanko et al., 2021a), yet the kinetic outcomes remain unknown and require further research. Third, our results do not fully represent kinetic analyses that can be performed completely outside the lab, which would also require a portable system to measure external forces addition to motion. Novel techniques such as force-sensing shoe insoles can be useful (Hullfish and Baxter, 2019). Regardless, our study is an important step towards this goal by providing the first evidence of 3D joints kinetics quantified with the portable markerless motion capture.

Markerless motion capture holds exciting potential to overcome many practical limitations of the conventional marker-based systems. The suitability of markerless motion capture for research and clinical use depends on its accuracy in quantifying the biomechanics of human motion. In this study, we found that markerless motion capture estimates of lower extremity kinematics and kinetics are highly comparable to marker-based in most measures. At the ankle and knee joints, where angle and moment waveforms match closely and magnitude differences are minimal during slower-speed movements, markerless motion capture can be used to facilitate large-scale and real-world biomechanical assessments not previously feasible with marker-based techniques. Between-system differences were larger during rapid movements and at the hip, indicating the need to compare against gold-standard measurements and establish best practices. The biomechanics community should continue to verify, validate, and expand the utility of markerless motion capture, thereby advancing collaborative research and clinical translation.

## Acknowledgments

Funding of this study was supported by the National Institutes of Health NIAMS R01AR078898, R01AR072034, and NICHD R37HD037985. The authors thank Madison Woods, Audrey Lehneis, and Liliann Zou for their assistance with data processing.

## Data statement

The motion capture system parameters, biomechanical models, and processed data presented in this article are available upon request to the corresponding (K.S.) or the senior author (J.R.B.).

## Supplementary material

Supplementary material of this study can be found in the online version of this article.

## References

1. Akbarshahi, M., Schache, A.G., Fernandez, J.W., Baker, R., Banks, S., Pandy, M.G., 2010. Non-invasive assessment of soft-tissue artifact and its effect on knee joint kinematics during functional activity. J. Biomech. 43, 1292–1301. doi: 10.1016/j.jbiomech.2010.01.002.

2. Andriacchi, T.P., Strickland, A.B., 1985. Gait analysis as a tool to assess joint kinetics. In: Berme, N., Engin, A.E., da Silva, K.M.C. (Eds.), Biomechanics of Normal and Pathological Human Articulating Joints. Springer, Dordrecht, Netherlands, pp. 83–102.

3. Camomilla, V., Cereatti, A., Cutti, A.G., Fantozzi, S., Stagni, R., Vannozzi, G., 2017. Methodological factors affecting joint moments estimation in clinical gait analysis: a systematic review. Biomed. Eng. Online. 16, 106. doi: 10.1186/s12938-017-0396-x.

4. Cappozzo, A., Della Croce, U., Leardini, A., Chiari, L., 2005. Human movement analysis using stereophotogrammetry: Part 1: theoretical background. Gait Posture 21, 186–196. doi: 10.1016/j.gaitpost.2004.01.010.

5. Chiari, L., Della Croce, U., Leardini, A., Cappozzo, A., 2005. Human movement analysis using stereophotogrammetry: Part 2: Instrumental errors. Gait Posture 21, 197–211. doi: 10.1016/j.gaitpost.2004.04.004.

6. Colyer, S.L., Evans, M., Cosker, D.P., Salo, A.I.T., 2018. A review of the evolution of vision-based motion analysis and the integration of advanced computer vision methods towards developing a markerless system. Sports Med. Open 4, 24. doi: 10.1186/s40798-018-0139-y.

7. Cronin N.J., 2021. Using deep neural networks for kinematic analysis: Challenges and opportunities. J. Biomech. 123, 110460. doi: 10.1016/j.jbiomech.2021.110460.

8. Della Croce, U., Leardini, A., Chiari, L., Cappozzo, A., 2005. Human movement analysis using stereophotogrammetry: Part 4: assessment of anatomical landmark misplacement and its effects on joint kinematics. Gait Posture 21, 226–237. doi: 10.1016/j.gaitpost.2004.05.003.

9. Derrick, T.R., van den Bogert, A.J., Cereatti, A., Dumas, R., Fantozzi, S., Leardini, A., 2020. ISB recommendations on the reporting of intersegmental forces and moments during human motion analysis. J. Biomech. 99, 109533. doi: 10.1016/j.jbiomech.2019.109533.

10. Drazan, J.F., Phillips, W.T., Seethapathi, N., Hullfish, T.J., Baxter, J.R., 2021. Moving outside the lab: Markerless motion capture accurately quantifies sagittal plane kinematics during the vertical jump. J. Biomech. 125, 110547. doi: 10.1016/j.jbiomech.2021.110547.

11. Hamner, S.R., Seth, A., Delp, S.L., 2010. Muscle contributions to propulsion and support during running. J. Biomech. 43, 2709–2716. doi: 10.1016/j.jbiomech.2010.06.025.

12. Hullfish, T.J., Baxter, J.R., 2020. A simple instrumented insole algorithm to estimate plantar flexion moments. Gait Posture 79, 92–95. doi: 10.1016/j.gaitpost.2020.04.016.

13. Ito, N., Sigurðsson, H.B., Seymore, K.D., Arhos, E.K., Buchanan, T.S., Snyder-Mackler, L., Silbernagel, K.G., 2022. Markerless motion capture: What clinician-scientists need to know right now. JSAMS Plus 1, 100001. doi: 10.1016/j.jsampl.2022.100001.

14. Kanko, R.M., Laende, E.K., Davis, E.M., Selbie, W.S., Deluzio, K.J., 2021a. Concurrent assessment of gait kinematics using marker-based and markerless motion capture. J. Biomech. 127, 110665. doi: 10.1016/j.jbiomech.2021.110665.

15. Kanko, R.M., Laende, E., Selbie, W.S., Deluzio, K.J., 2021b. Inter-session repeatability of markerless motion capture gait kinematics. J. Biomech. 121, 110422. doi:10.1016/j.jbiomech.2021.110422.

16. Kanko, R.M., Laende, E.K., Strutzenberger, G., Brown, M., Selbie, W.S., DePaul, V., Scott, S.H., Deluzio, K.J., 2021c. Assessment of spatiotemporal gait parameters using a deep learning algorithm-based markerless motion capture system. J. Biomech. 122, 110414. doi:10.1016/j.jbiomech.2021.110414.

17. Kapron, A.L., Aoki, S.K., Peters, C.L., Maas, S.A., Bey, M.J., Zauel, R., Anderson, A.E., 2014. Accuracy and feasibility of dual fluoroscopy and model-based tracking to quantify in vivo hip kinematics during clinical exams. J. Appl. Biomech. 30, 461–470. doi: 10.1123/jab.2013-0112.

18. Keller, V.T., Outerleys, J.B., Kanko, R.M., Laende, E.K., Deluzio, K.J., 2022. Clothing condition does not affect meaningful clinical interpretation in markerless motion capture. J. Biomech. 141, 111182. doi: 10.1016/j.jbiomech.2022.111182.

19. Leardini, A., Chiari, L., Della Croce, U., Cappozzo, A., 2005. Human movement analysis using stereophotogrammetry: Part 3. Soft tissue artifact assessment and compensation. Gait Posture 21, 212–225. doi: 10.1016/j.gaitpost.2004.05.002.

20. Li, G., Wuerz, T.H., DeFrate, L.E., 2004. Feasibility of using orthogonal fluoroscopic images to measure in vivo joint kinematics. J. Biomech. Eng. 126, 314–318. doi: 10.1115/1.1691448.

21. Mathis, A., Mamidanna, P., Cury, K.M., Abe, T., Murthy, V.N., Mathis, M.W., Bethge, M., 2018. DeepLabCut: markerless pose estimation of user-defined body parts with deep learning. Nat Neurosci. 21, 1281–1289. doi: 10.1038/s41593-018-0209-y.

22. Moisio, K.C., Sumner, D.R., Shott, S., Hurwitz, D.E., 2003. Normalization of joint moments during gait: a comparison of two techniques. J. Biomech. 36, 599–603. doi: 10.1016/s0021-9290(02)00433-5.

23. Nester, C., Jones, R.K., Liu, A., Howard, D., Lundberg, A., Arndt, A., Lundgren, P., Stacoff, A., Wolf, P., 2007. Foot kinematics during walking measured using bone and surface mounted markers. J. Biomech. 40, 3412–3423. doi: 10.1016/j.jbiomech.2007.05.019.

24. Novacheck T.F., 1998. The biomechanics of running. Gait Posture 7, 77–95. doi: 10.1016/s0966-6362(97)00038-6.

25. Riley, P.O., Dicharry, J., Franz, J., Della Croce, U., Wilder, R.P., Kerrigan, D.C., 2008. A kinematics and kinetic comparison of overground and treadmill running. Med. Sci. Sports Exerc. 40, 1093–1100. doi: 10.1249/MSS.0b013e3181677530.

26. Robertson, D.G.E., Caldwell, G.E., Hamill, J., Kamen, G., Whittlesey, S. (Eds.), 2013. Research Methods in Biomechanics. Human Kinetics, Champaign, IL.

27. Schober, P., Boer, C., Schwarte, L.A., 2018. Correlation coefficients: appropriate use and interpretation. Anesth. Analg. 126, 1763–1768. doi: 10.1213/ANE.0000000000002864.

28. Slater, A.A., Hullfish, T.J., Baxter, J.R., 2018. The impact of thigh and shank marker quantity on lower extremity kinematics using a constrained model. BMC Musculoskelet. Disord. 19, 399. doi: 10.1186/s12891-018-2329-7.

29. Stagni, R., Leardini, A., Cappozzo, A., Benedetti, M.G., Cappello, A., 2000. Effects of hip joint centre mislocation on gait analysis results. J. Biomech. 33, 1479–1487. doi: 10.1016/s0021-9290(00)00093-2.

30. Tang, H., Pan, J., Munkasy, B., Duffy, K., Li, L., 2022. Comparison of lower extremity joint moment and power estimated by markerless and marker-based systems during treadmill running. Bioengineering (Basel) 9, 574. doi: 10.3390/bioengineering9100574.

31. Tashman, S., Anderst, W., 2003. In-vivo measurement of dynamic joint motion using high speed biplane radiography and CT: application to canine ACL deficiency. J. Biomech. Eng. 125, 238–245. doi: 10.1115/1.1559896.

32. Wu, G., Siegler, S., Allard, P., Kirtley, C., Leardini, A., Rosenbaum, D., Whittle, M., D’Lima, D.D., Cristofolini, L., Witte, H., Schmid, O., Stokes, I., 2002. ISB recommendation on definitions of joint coordinate system of various joints for the reporting of human joint motion – part I: ankle, hip, and spine. J. Biomech. 35, 543–548. doi: 10.1016/s0021-9290(01)00222-6.

33. Wu, G., van der Helm, F.C., Veeger, H.E., Makhsous, M., Van Roy, P., Anglin, C., Nagels, J., Karduna, A.R., McQuade, K., Wang, X., Werner, F.W., Buchholz, B., 2005. ISB recommendation on definitions of joint coordinate systems of various joints for the reporting of human joint motion – Part II: shoulder, elbow, wrist and hand. J. Biomech. 38, 981–992. doi: 10.1016/j.jbiomech.2004.05.042.

