## Supplementary Material 1 for "Markerless motion capture estimates of lower extremity kinematics and kinetics are comparable to marker-based across 8 movements"

### Supplementary material 1. Lower extremity kinematics and kinetics estimated from markerless motion capture compared to marker-based across a total of 36 movements

The secondary data presented herein are extended from the same biomechanical analysis we performed on the 8 movements presented in the main article. Each subject performed all 36 movements during the experiment of a larger study, in the order identical to the result Figures presented below (i.e., heel raises, walking, etc…) including the 8 representative movements (highlighted title in teal color). Waveforms = group mean (line) ± 1 SD (shade) for marker-based (blue), markerless (red), and their difference (gold).

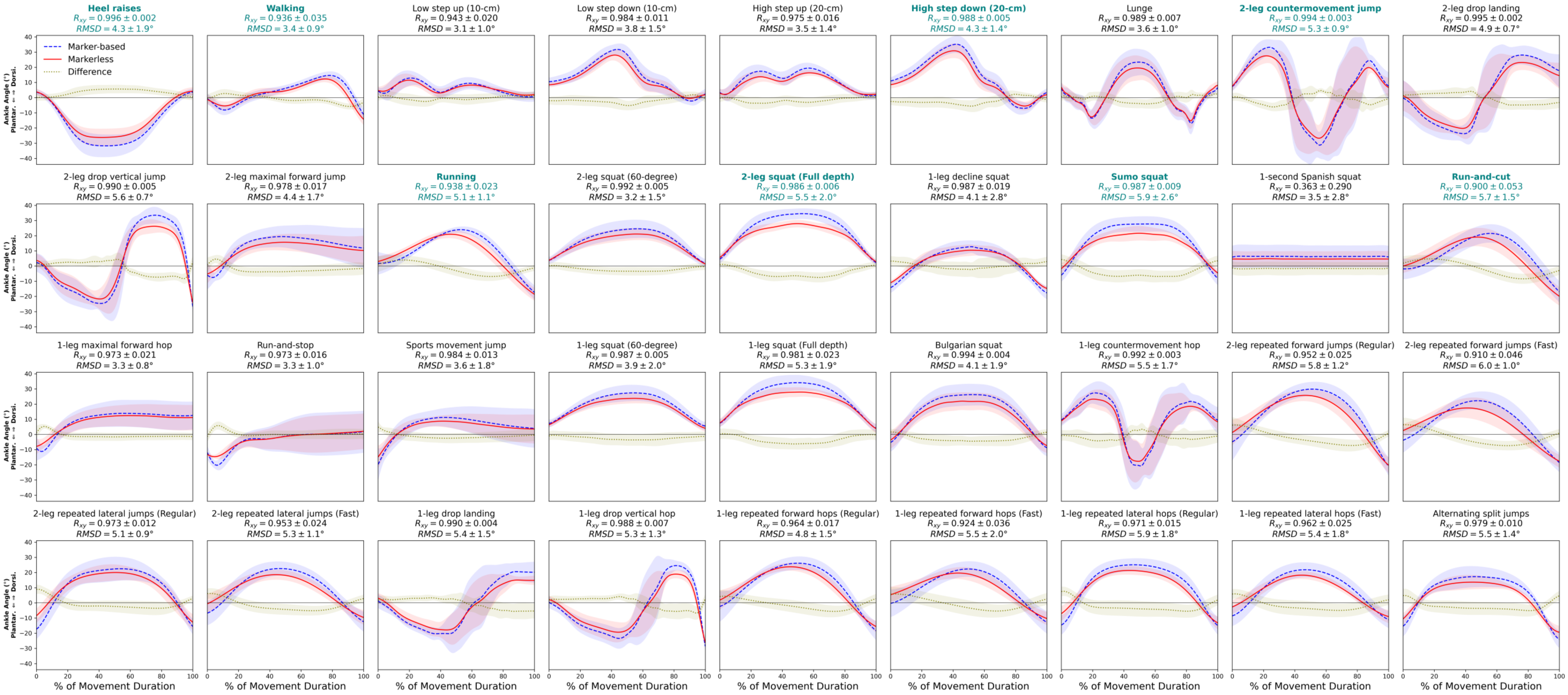


**Supplementary Figure 1.** Kinematics: ankle dorsiflexion (+) / plantarflexion (-) angle.

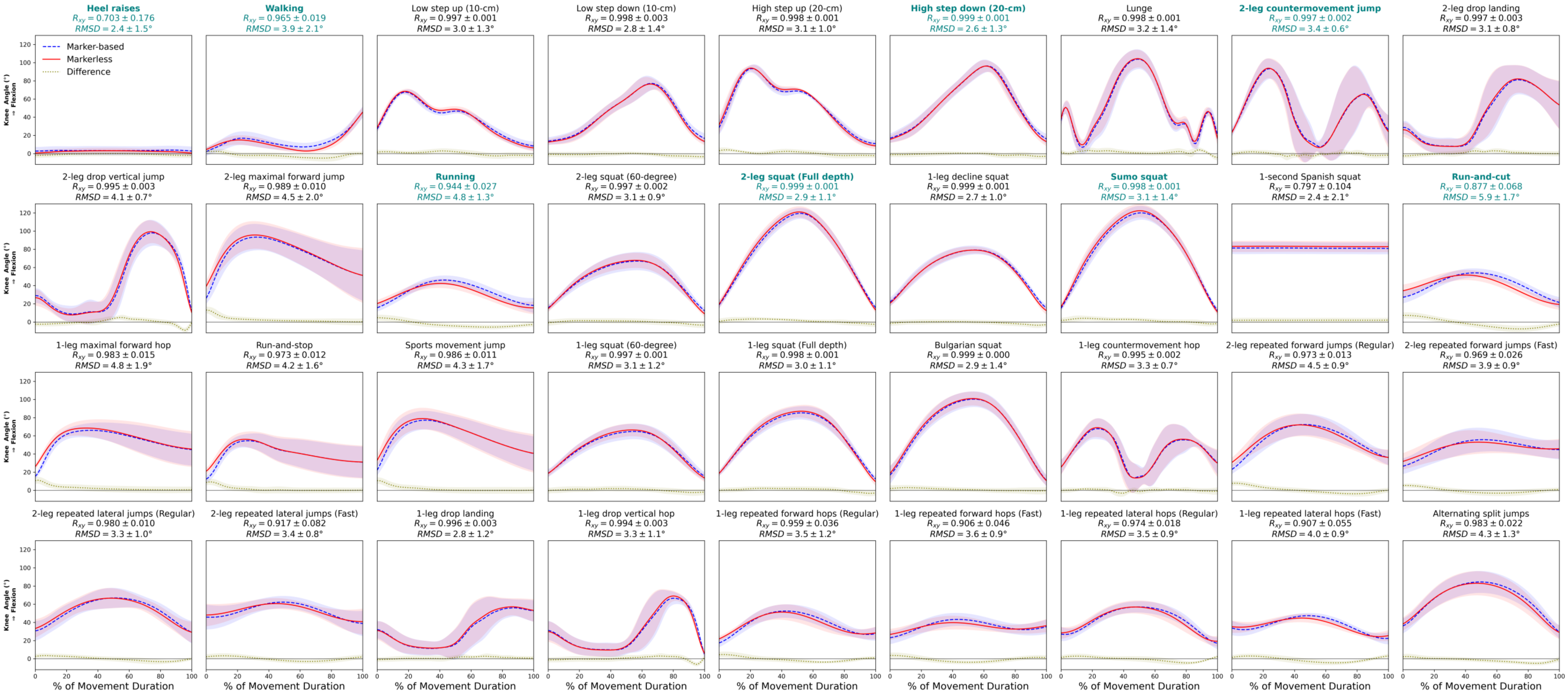


**Supplementary Figure 2.** Kinematics: knee flexion (+) angle.

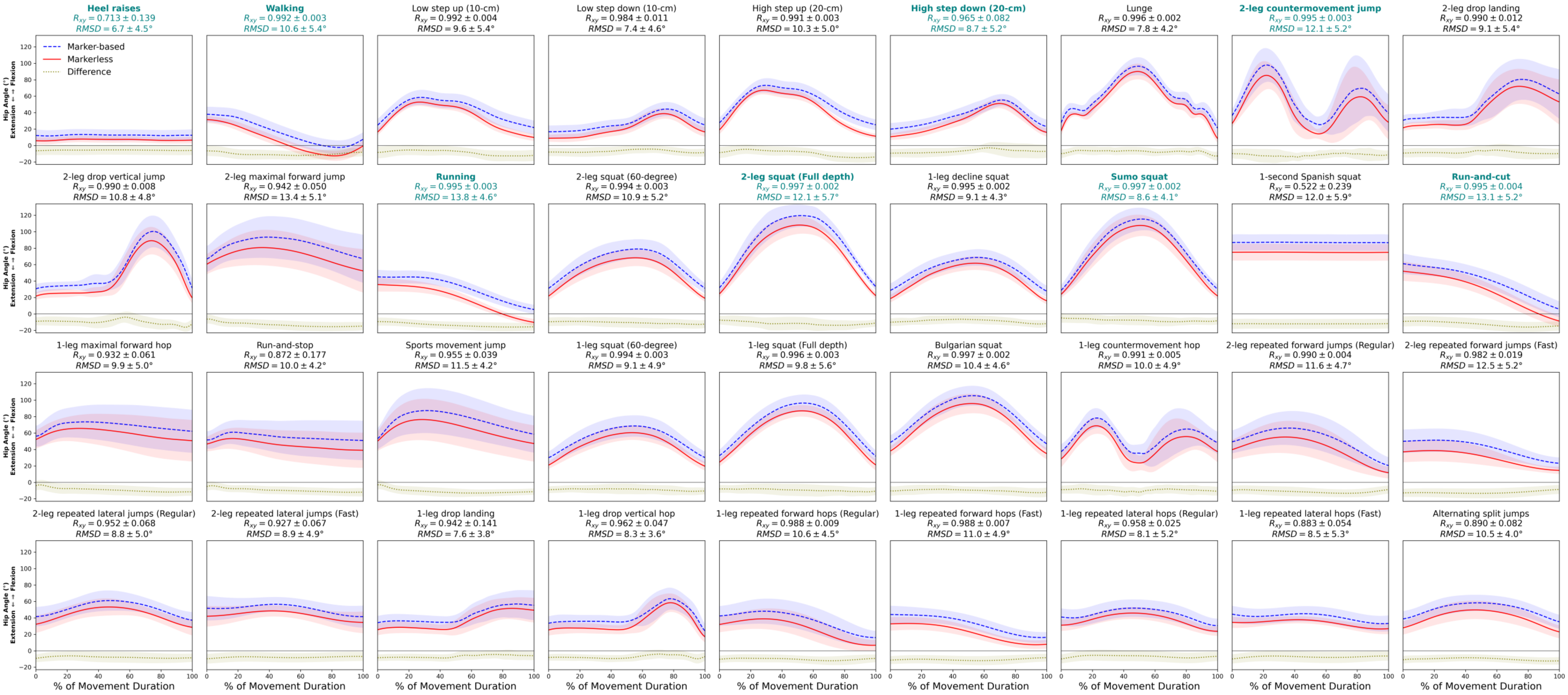


**Supplementary Figure 3.** Kinematics: hip flexion (+) / extension (-) angle.

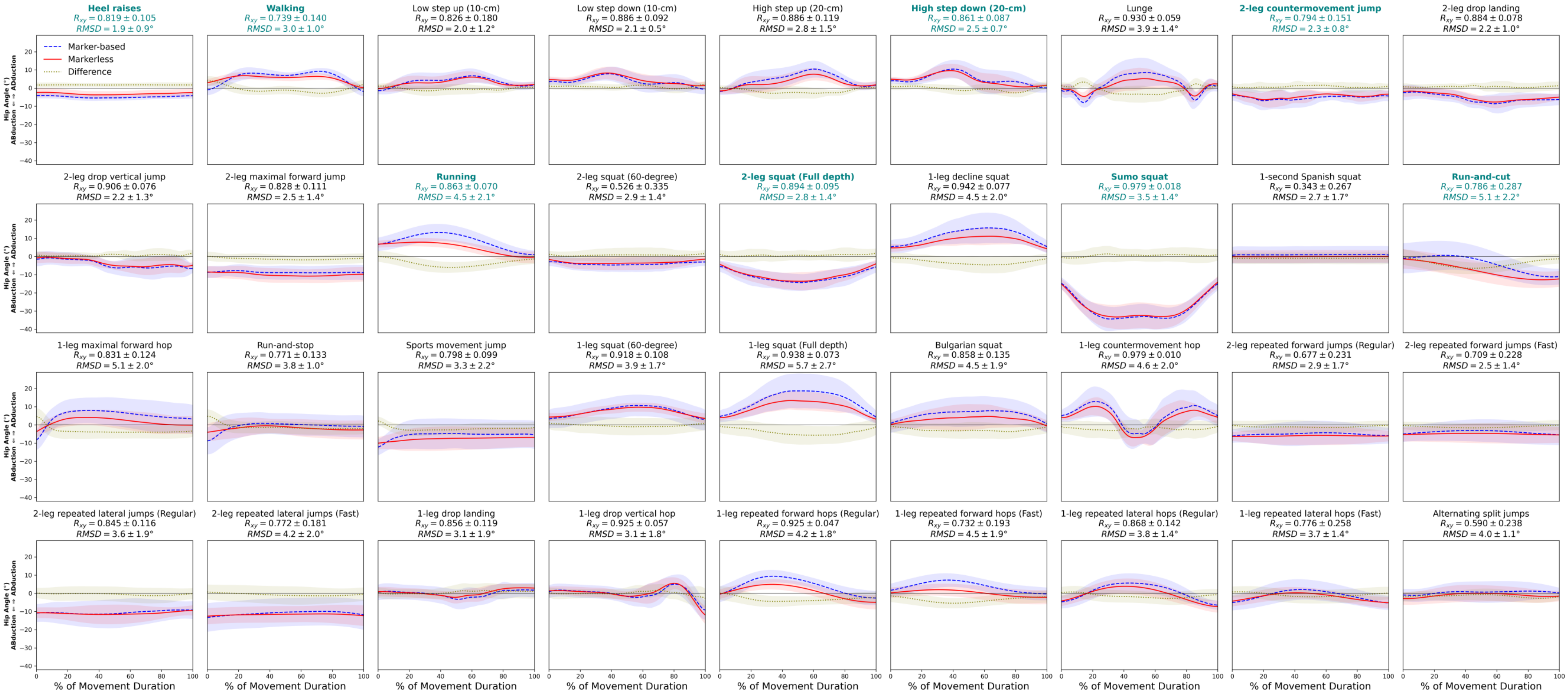


**Supplementary Figure 4.** Kinematics: hip adduction (+) / abduction (-) angle.

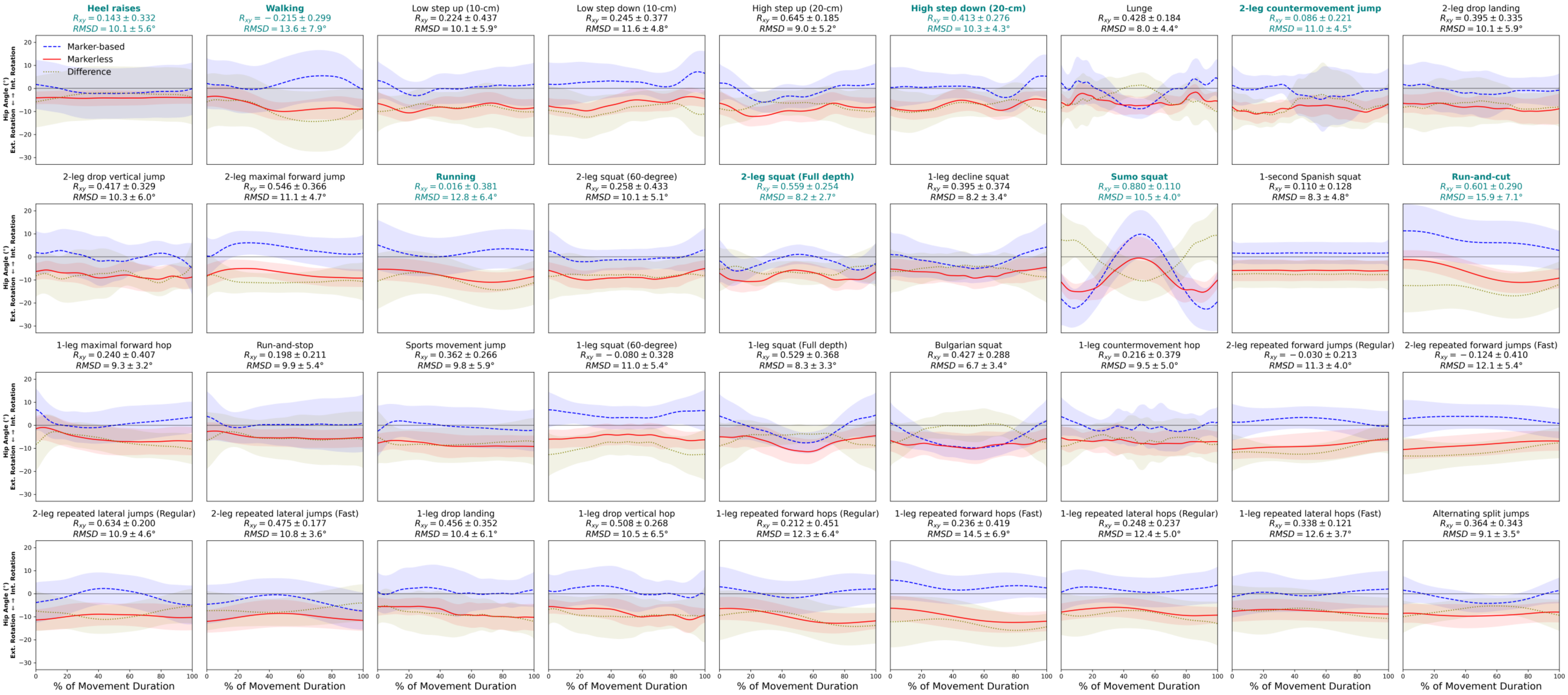


**Supplementary Figure 5.** Kinematics: hip internal (+) / external (-) rotation angle.

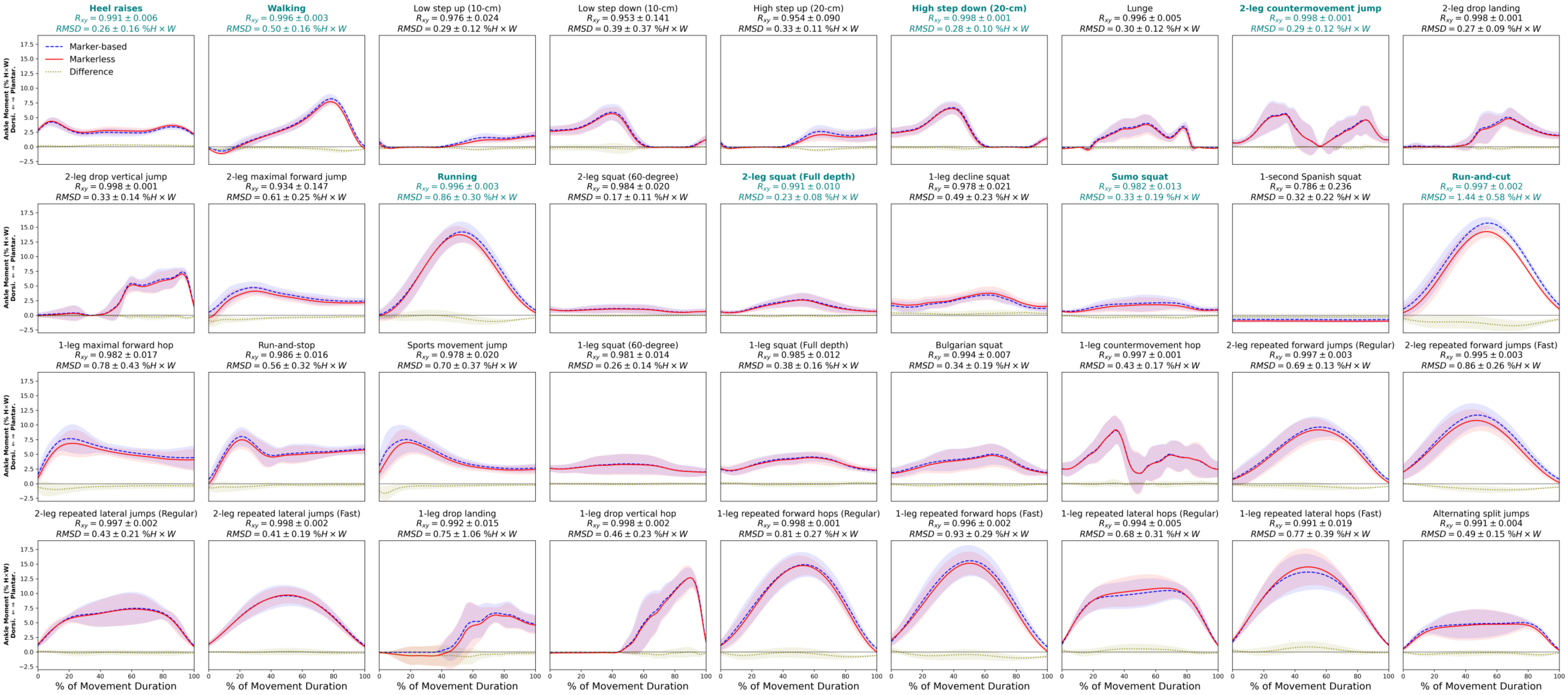


**Supplementary Figure 6.** Kinetics: ankle plantarflexion (+) / dorsiflexion (-) moment.

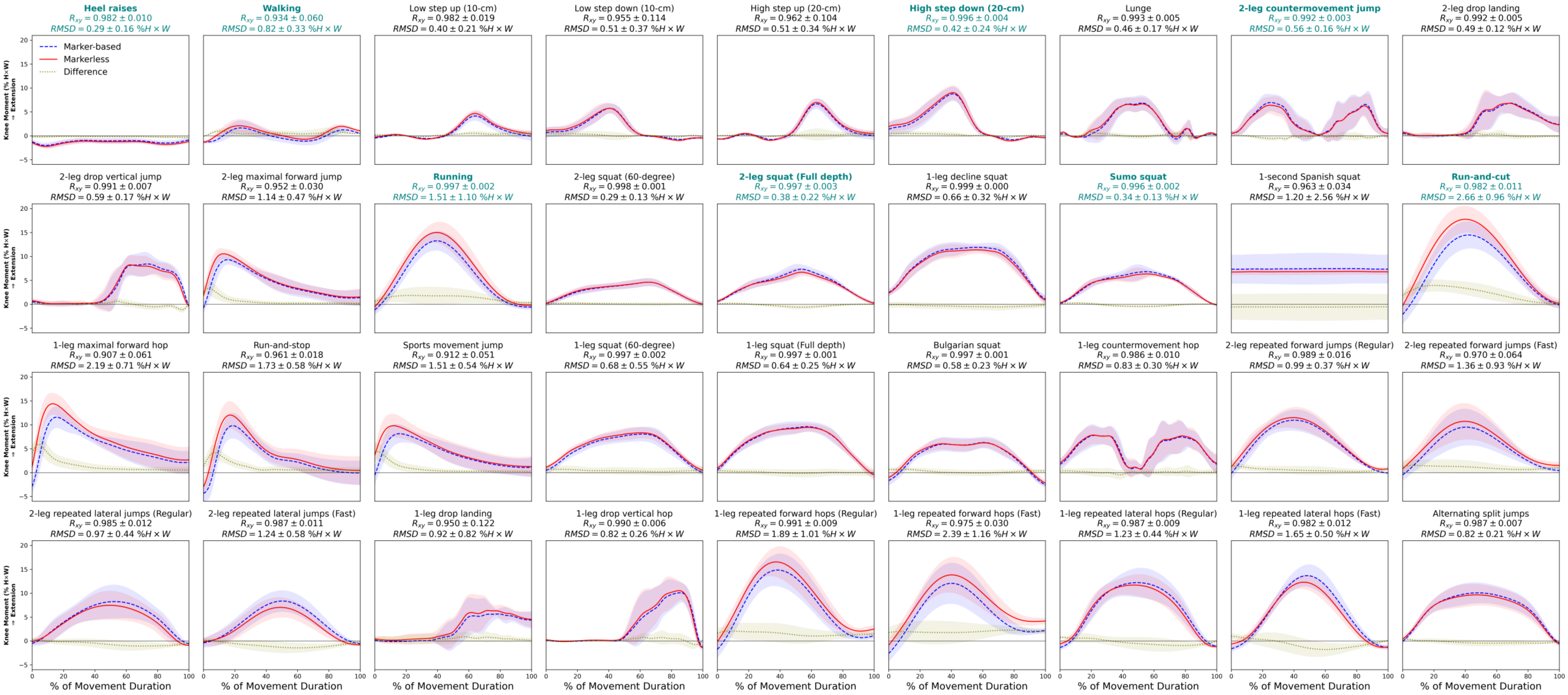


**Supplementary Figure 7.** Kinetics: knee extension (+) moment.

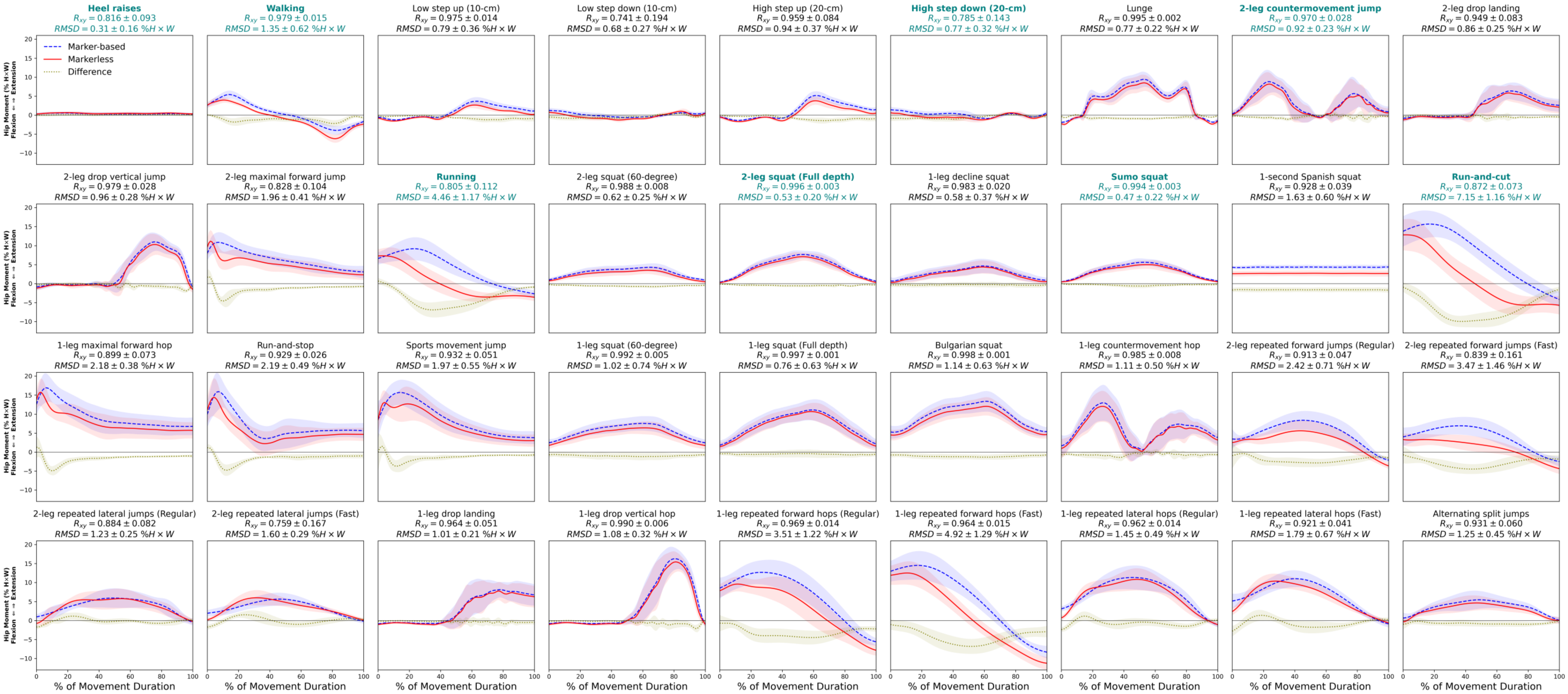


**Supplementary Figure 8.** Kinetics: hip extension (+) / flexion (-) moment.

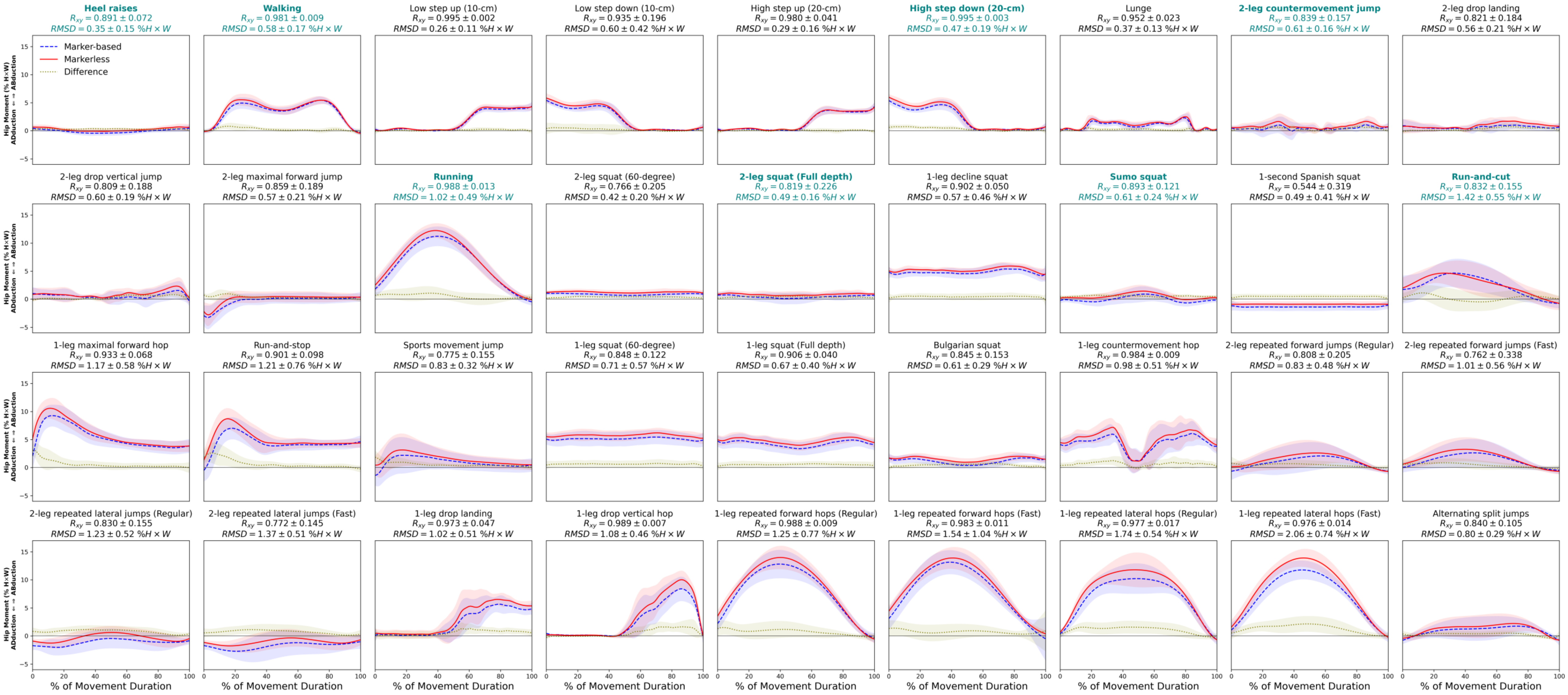


**Supplementary Figure 9.** Kinetics: hip abduction (+) / adduction (-) moment.

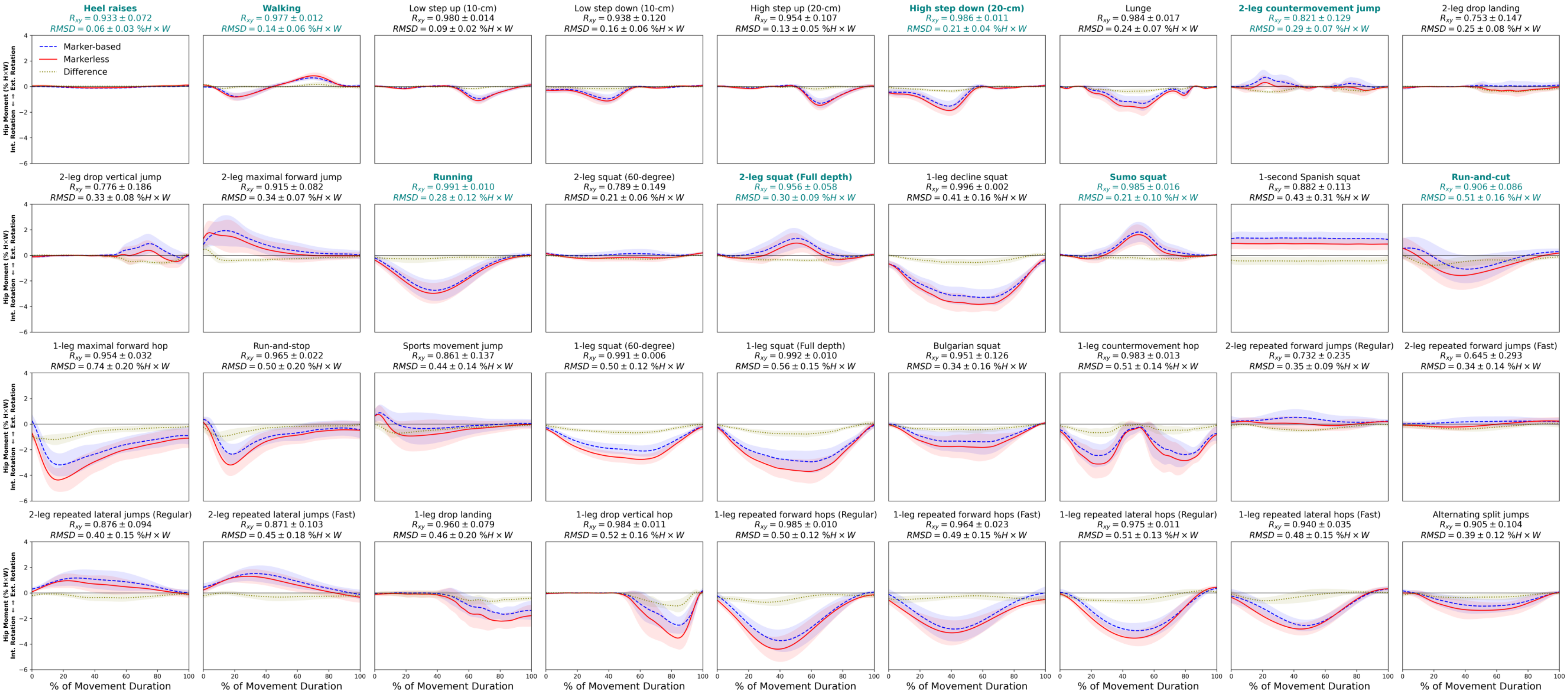


**Supplementary Figure 10.** Kinetics: hip external (+) / internal (-) rotation moment.

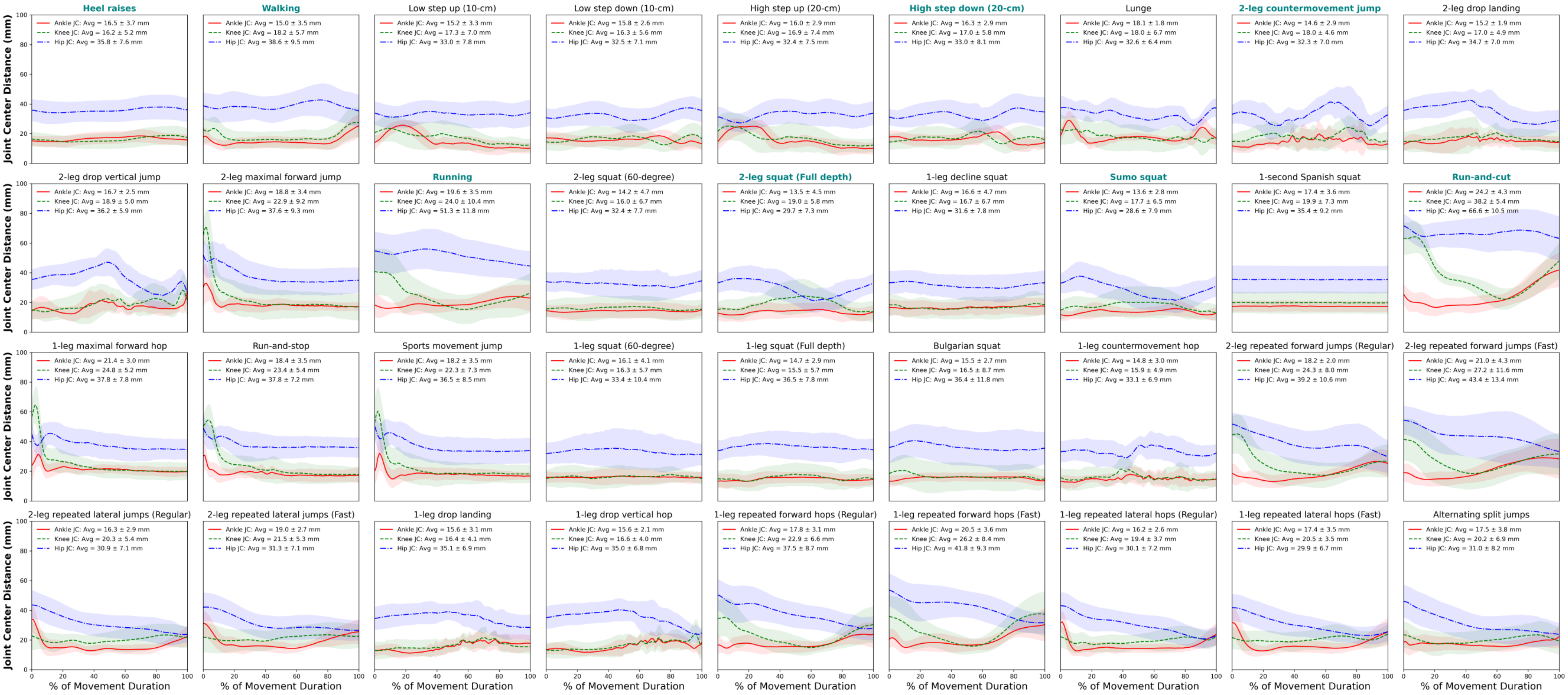


**Supplementary Figure 11.** Distance between the two joint centers tracked by either system.
