## Supplementary Material 2 for "Markerless motion capture estimates of lower extremity kinematics and kinetics are comparable to marker-based across 8 movements"

**Supplementary material 2. Joint center location differences between marker-based and markerless motion capture across 8 movements**

We calculated the right ankle, knee, and hip joint center locations estimated from both systems as a secondary measure that theoretically determines external force moment arms and affects joint kinetics (Robertson et al., 2013). We quantified these joint center locations during individual repetitions within each movement trial. In the marker-based model, we defined ankle joint as the midpoint between the lateral and medial malleoli anatomic landmarks, knee joint as the midpoint between the lateral and medial femoral epicondyles, and hip joint using regression equation distances offset from the anterior and posterior superior iliac spines (Harrington et al., 2007). In the Theia3D markerless model, the three joint center locations were estimated directly as human salient features from the deep learning algorithm (Kanko et al., 2021).

We computed the resultant Euclidian distance between the two locations of each joint, one tracked by the marker-based system and the other by the markerless. We then computed the time-wise mean of this resultant distance, which yielded 3 joint center distances (ankle, knee, hip) for each repetition. We averaged each variable across repetitions within a trial, then calculated the group mean of these variables across the 10 subjects to determine an overall level of joint location disagreement between the marker-based and markerless systems.

***Results.*** Ankle and knee joint center distances between marker-based and markerless motion capture were both small and generally consistent across the 8 movements, while the hip joint center distances between the two systems were larger especially during running and run-and-cut (**Suppl. Figure**). Among the other 6 relatively slower movements (i.e., except running and run-and-cut), the mean joint center distances ranged 13.5 – 16.5 mm at the ankle, 16.2 – 19.0 mm at the knee, and 28.6 – 38.6 mm at the hip. In contrast, joint center distances increased to 19.6 mm at the ankle, 24.0 mm at the knee, 51.3 mm at the hip during running (**Suppl. Figure d**), and 24.2 mm at the ankle, 38.2 mm at the knee, 66.6 mm at the hip during run-and-cut (**Suppl. Figure h**). For these two rapid movements, joint center location differences were relatively larger during early deceleration phases at the knee, and throughout the course of movement duration at the hip (**Suppl. Figure d, h**).

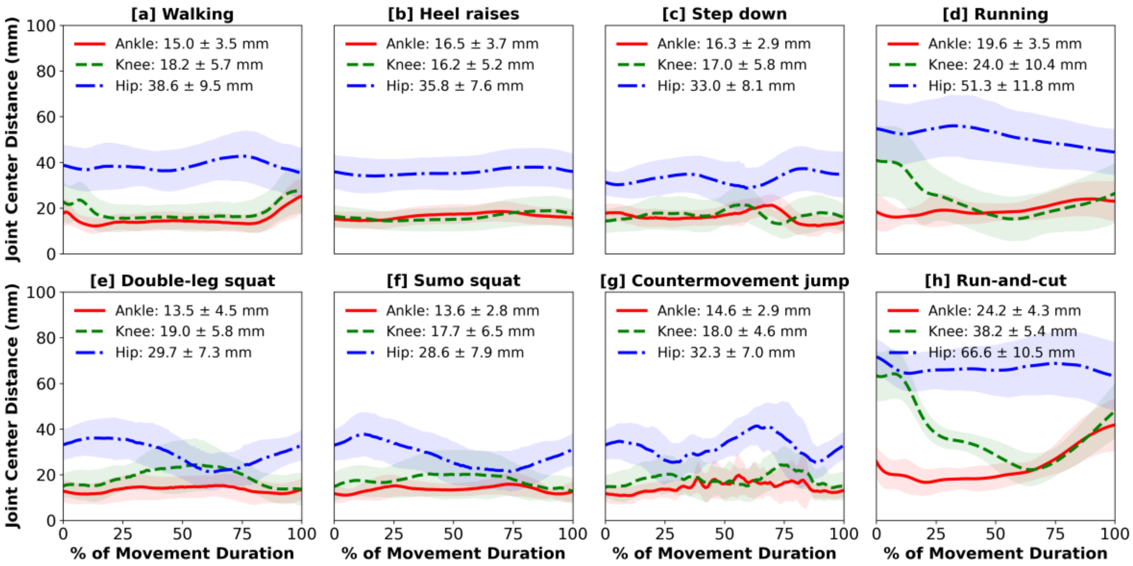


**Supplementary Figure.** Resultant distance between the two ankle (red), knee (green), and hip (blue) joint centers tracked by either system during 8 movements: [**a**] walking, [**b**] heel raises, [**c**] step-down, [**d**] running, [**e**] double-leg squat, [**f**] sumo squat, [**g**] countermovement jump, [**h**] run-and-cut. Waveforms = group mean (line) ± 1 SD (shade).

***Discussion.*** The inter-system joint location differences we found during walking were comparable to the results by Kanko et al. (2021). We also found that joint center locations were most different between the two systems at the hip, especially during running and run-and-cut

(**Suppl. Figure d, h**), which matched the joint moment differences (**Figure 4d, 5h**, main article). During running, Skin artifacts around the hip can be large in the sagittal plane, while ground strike also induces artifacts at the knee. Because the marker-based system relies on skin markers to approximate the joint landmarks underneath, increased skin motion would confound joint center tracking, which directly alter moment arms and propagate to joint kinetic errors. This may explain why we saw the largest kinetic differences in hip extension moments and early-phase knee extension moments (**Figure 4d, 5h**, main article). When assessing running motion, the skin artifact-free markerless motion capture can likely provide more accurate joint tracking.

### References


1. Harrington, M.E., Zavatsky, A.B., Lawson, S.E., Yuan, Z., Theologis, T.N., 2007. Prediction of the hip joint centre in adults, children, and patients with cerebral palsy based on magnetic resonance imaging. J. Biomech. 40, 595–602. doi:10.1016/j.jbiomech.2006.02.003
2. Kanko, R.M., Laende, E.K., Davis, E.M., Selbie, W.S., Deluzio, K.J., 2021a. Concurrent assessment of gait kinematics using marker-based and markerless motion capture. J. Biomech. 127, 110665. doi: 10.1016/j.jbiomech.2021.110665.
3. Robertson, D.G.E., Caldwell, G.E., Hamill, J., Kamen, G., Whittlesey, S. (Eds.), 2013. Research Methods in Biomechanics. Human Kinetics, Champaign, IL.
